# Differential expression analysis of log-ratio transformed counts: benchmarking methods for RNA-Seq data

**DOI:** 10.1101/231175

**Authors:** Thomas P. Quinn, Tamsyn M. Crowley, Mark F. Richardson

## Abstract

**Background:** Count data generated by next-generation sequencing assays do not measure absolute transcript abundances. Instead, the data are constrained to an arbitrary “library size” by the sequencing depth of the assay, and typically must be normalized prior to statistical analysis. The constrained nature of these data means one could alternatively use a log-ratio transformation in lieu of normalization, as often done when testing for differential abundance (DA) of operational taxonomic units (OTUs) in 16S rRNA data. Therefore, we benchmark how well the **ALDEx2** package, a transformation-based DA tool, detects differential expression in high-throughput RNA-sequencing data (RNA-Seq), compared to conventional RNA-Seq differential expression methods.

**Results:** To evaluate the performance of log-ratio transformation-based tools, we apply the **ALDEx2** package to two simulated, and one real, RNA-Seq data sets. The latter was previously used to benchmark dozens of conventional RNA-Seq differential expression methods, enabling us to directly compare transformation-based approaches. We show that **ALDEx2**, widely used in meta-genomics research, identifies differentially expressed genes (and transcripts) from RNA-Seq data with high precision and, given sufficient sample sizes, high recall too (regardless of the alignment and quantification procedure used). Although we show that the choice in log-ratio transformation can affect performance, **ALDEx2** has high precision (i.e., few false positives) across all transformations. Finally, we present a novel, iterative log-ratio transformation (now implemented in **ALDEx2**) that further improves performance in simulations.

**Conclusions:** Our results suggest that log-ratio transformation-based methods can work to measure differential expression from RNA-Seq data, provided that certain assumptions are met. Moreover, these methods have high precision (i.e., few false positives) in simulations and perform as good as, or better than, than conventional methods on real data. With previously demonstrated applicability to 16S rRNA data, **ALDEx2** can work as a single tool for data from multiple sequencing modalities.

## Background

In the last decade, new technologies, collectively known as next generation sequencing (NGS), have come to dominate the market [22]. Although NGS has a wide range of applications, the use of NGS in transcriptome profiling, called massively parallel RNA-sequencing (RNA-Seq), is perhaps most popular [41]. Like microarray, RNA-Seq is used to quantify transcript abundance (i.e., expression) [22]. Unlike microarrays however, RNA-Seq is able to estimate the abundance of uncharacterized transcripts as well as differentiate between transcript isoforms [41]. Meanwhile, advances in NGS have reduced the cost of sequencing tremendously, making it possible to generate an enormous amount of raw sequencing data easily and cheaply.

However, the analysis of raw sequencing data is not trivial. The data, constituting a “library” of hundreds of thousands of short sequence fragments, must undergo a number of processing steps prior to abundance estimation [15]. In the setting of an established reference genome (or transcriptome), this process generally includes (1) removal of undesired sequences (e.g., assay-specific adapters, ribosomal RNA, or short reads) and quality filtering, (2) alignment of the remaining sequences to the reference, and (3) quantification of transcript abundance [15]. In addition, RNA-Seq has two notable sources of bias that an analyst may need to address: transcripts with longer lengths [23] and higher GC content [9] have their abundances over-estimated.

Alignment is a computationally expensive process that, in many cases, contributes to the major bottleneck in RNA-Seq workflows [8]. As the ability to generate raw sequence data appears to outpace gains in computing power, the advantage of fast alignment seems clear. Although dozens of aligners exist (e.g., see [16] [4] [40] [3]), the **STAR** aligner [8] has grown in popularity as a method that balances accuracy with efficiency, having good performance in systematic evaluations [10] [3]. Recently, a new family of “pseudo-alignment” methods has emerged (e.g., `Kallisto` [6], **`Sailfish`** [26], and **`Salmon`** [25]), providing an order of magnitude faster speeds than conventional aligners [25]. On the other hand, quantification is comparatively quick and usually performed by the aligners themselves. In essence, quantification involves “counting” the number of times a sequence aligns to a given portion of the reference [15]. This results in a matrix of counts (or pseudo-counts) describing the estimated number of times each transcript was present for each sample under study (although some methods represent abundance in other units [37]). Yet, the choice in the alignment and quantification method used seems to matter less than the choice in the software used for down-stream analyses[43].

The “count matrix” produced by alignment and quantification is perhaps most commonly used for differential expression (DE) analysis, a means by which to identify which genes (or transcripts), if any, have a statistically significant difference in abundance across the experimental groups [15]. Like alignment, dozens of methods exist for DE analysis, providing a unique approach to normalization and statistical modeling. Of these, **`DESeq`** [2] and `edgeR` [30] seem most popular. Both use a type of normalization whereby each “library” (i.e., sample vector) is adjusted by a scaling factor based on a reference (or pseudo-reference). **`DESeq`** uses as the reference the median of the ratios of each gene for that sample to the geometric mean of each gene for all samples [2] [7]. Meanwhile, **`edgeR`** uses as the reference the weighted mean of log ratios between that sample and an explicitly chosen reference, a method known as the trimmed mean of M (TMM) [31] [7]. Underlying this approach is the rarely stated assumption that most transcripts do not differ in abundance while gains and losses happen with equipoise [11].

While the choice in normalization can affect the final results of a DE analysis [19] [18], it is necessary because per-sample counts generated by alignment and quantification do not compare directly [33]. This is because a sequencer only sequences a fraction of the total input, thereby constraining the per-sample output to a fraction of the total amount of sequencing (called sequencing depth) [33]. As such, the increased presence of any one transcript in the input material results in a decreased measurement for all other transcripts [33]. This sum constraint makes RNA-Seq data a kind of compositional data in which each sample is a composition and the transcript-wise counts are the components [20]. Compositional data have two key properties. First, the total sum of all components is an artifact of the sampling procedure [39]. Second, the differences between any two components only carry meaning proportionally [39]. For example, the difference between the two counts [50, 500] is the same as the difference between [100, 1000] since the latter could be obtained from the former simply by doubling the sequencing depth. RNA-Seq data have both of these properties, but differ slightly from true compositional data in that count data only contain integer values [20] [28].

Compositional data analysis (CoDA) describes a collection of methods used to analyze compositional data, including those pioneered by Aitchison in 1986 [1]. Commonly, such analyses begin with a transformation, most often the centered log-ratio (clr) transformation (defined in Methods). In contrast to normalizations, these transformations do not claim to retrieve absolute abundances from the compositional data [27]. Yet, they are sometimes used as if they were normalizations themselves [27]. The **ALDEx2** package (available for the R programming language) uses log-ratio transformations in lieu of normalization for the analysis of sequencing data [12]. This package, first developed for analyzing meta-genomics data, identifies differentially abundant features across two or more groups by applying statistical hypothesis testing to compositional data in three steps: (1) generate Monte Carlo (MC) instances (of log-ratio transformed data) based on the provided count matrix using the Dirichlet distribution, (2) apply univariate statistical models on the MC instances, and (3) calculate the expected (i.e., average) FDR-adjusted per-transcript *p*-values across all MC instances [12]. By default, **ALDEx2** uses the clr transformation, although it supports other transformations too.

The **ALDEx2** package is not well-adopted for RNA-Seq analysis, although its applicability to RNA-Seq is established elsewhere [13]. However, **ALDEx2** is used to analyze 16S rRNA data (e.g., [38] [5]) where it is shown to achieve much lower false positive rates (FPR) than competing DE methods [36]. Yet, we do not know of any paper which independently benchmarks **ALDEx2** as a DE method for RNA-Seq (excepting the aforementioned article written by the **ALDEx2** authors [13]). Moreover, we do not know the extent to which the choice of log-ratio transformation influences the final results of an analysis. Finally, we do not know whether the results produced by **ALDEx2** are sensitive to the chosen alignment and quantification method. In this paper, we use simulated and real data to evaluate the performance of **ALDEx2** as a DE method for RNA-Seq data and demonstrate that **ALDEx2**, under the appropriate conditions, performs as well as or better than standard DE methods, establishing its validity for the analysis of RNA-Seq data. In doing so, we also present a novel log-ratio transformation, based on iterative runs of **ALDEx2**, that improves accuracy when compared with other approaches.

## Methods

### Data acquisition

To benchmark how well the **ALDEx2** package (available for the R programming language) performs as a differential expression method for RNA-Seq data, we analyzed three data sets. The first two contain simulated data generated from the polyester package (available for the R programming language) [14]. polyester simulates RNA-Seq data as raw sequencing data (i.e., FASTQ read sets) where the abundances of the transcripts follow a negative binomial model [14]. The third data set contained real data sourced from a previously published benchmark study [43], retrieved as raw RNA sequencing data (i.e., FASTQ read sets) from the NCBI SRA, under accession SRP082682 https://www.ncbi.nlm.nih.gov/Traces/study/?acc=srp082682. We used these raw data for sequencing alignment, quantification, and differential expression analysis.

### Data simulation

We simulated two sequencing experiments with two groups of twenty samples each (i.e., 40 samples total per experiment) using a forked version of polyester, hosted on GitHub, that adds multi-core support (archived at https://github.com/kcha/polyester/commits/545e33c9776db2927f9a22c8c2f5bfde2b3081a7). As such, when describing data sets with less than 40 samples, we refer to a random sub-sample of the complete data set and not to newly simulated data.

To run polyester, we used the human **`GRCh37`** DNA primary assembly FASTA file and the **`GRCh37.87`** annotation GTF file, as compiled into a FA file using the gffread command line tool [37]. We set the parameters to achieve 20x coverage with a mean fragment length of 300 bases. Transcripts were selected randomly to have different magnitudes of differential expression with weighted probability: 4-fold up-regulation (3% of transcripts), 2-fold up (7%), 1.5-fold up (9%), 1.5-fold down-regulation (6%), 2-fold down (3%), and 4-fold down (2%). Each sample had a random multiplicative weight applied to their library size (with a mean between-group difference of 0.20 with a within-group standard deviation of 0.05).

Otherwise, the two simulated experiments differ only in the mean-variance relationship underlying the negative binomial model. The first is a low-variance data set built using the default **`size`** argument. The second is a high-variance data set built using **`size`** = 1 such that the variance of the negative binomial model equals the mean plus the mean squared. We selected these size parameters based on the precedent set by the authors in their flagship publication [14].

### Alignment and quantification

To maintain benchmarking comparisons with Williams et al. [43], we used alignment and quantification protocols (for each of our three data sets) congruent with theirs. We performed alignment to the **`GRCh37`** release of the Human genome using **`STAR`** v2.5 [8] and Salmon v0.8 [25]. For **`STAR`**, we used the “Basic” two-pass mode to output BAM alignments to the Human genome and transcriptome. For Salmon, alignments were made to the transcriptome (built using gffread, as for the simulated data) across 100 bootstraps. For Salmon, we trialled both the SMEM-based lightweight-alignment approach (hereafter called slFMD) and the “quasi-mapping” approach (hereafter called slQUASI).

We quantified expression at both the gene- and transcript-level. For transcript-level expression, counts were estimated using **`salmon quant`** for the slFMD and slQUASI alignments, as well as for the **`STAR`** transcriptome alignment (hereafter called stsl). For gene-level expression, counts were estimated using the “GeneCounts” quant mode in **`STAR`** (i.e., using the **`STAR`** alignment; hereafter called stst). We then condensed transcript-level expression to gene-level expression using the tximport package (available for the R programming language) [34] with the argument type = "salmon” and a key built from the EnsDb.Hsapiens.v86 database (from Bioconductor) [29].

### Log-ratio transformations

The **ALDEx2** package produces different results depending on the log-ratio transformation used. Two transformations available in **ALDEx2** are the centered log-ratio (clr) [1] and inter-quartile log-ratio (iqlr) transformations [12]. For *D* genes (or transcripts), the clr-transformation is defined as the logarithm of the transcript counts for the i-th sample, x, divided by the geometric mean of all counts for that sample, *g*(x):

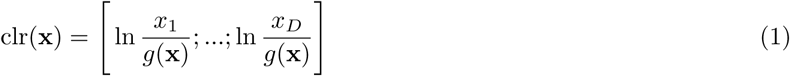

The iqlr-transformation replaces the *g*(x) denominator term with the geometric mean of those transcripts within the inter-quartile range of variability (i.e., prior to transformation), *g*(x_iqr_):

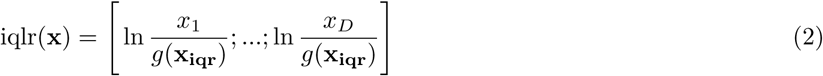

In the analysis of the simulated data, we also use what we call the multi-additive log-ratio (malr) transformation. This uses the identity of all equally expressed transcripts as a reference set. Although this transformation is only feasible here because we already know *a priori* which transcripts are differentially expressed, it provides a “best case scenario” for normalization against which to compare other transformations. This transformation replaces the *g*(x) denominator term with the geometric mean of equally expressed transcripts, *g*(x_eq_):

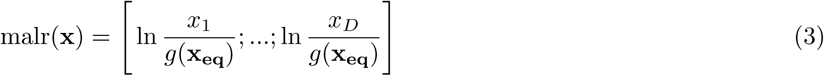

In addition, we introduce a novel transformation called the iterative iqlr (iilr) transformation. The iilr-transformation begins with the familiar iqlr-transformation, but then uses the results of a complete **ALDEx2** analysis to inform a subsequent iteration of **ALDEx2**. After the initial iqlr-transformation, each new **ALDEx2** run uses the geometric mean of the equally expressed transcripts identified by the prior **ALDEx2** run. In principle, the equally expressed transcripts identified by each new iteration of **ALDEx2** should more closely approximate the idealized x_eq_ used by the malr. We trial a single iteration of the iilr-transformation (ii1) as well as an approach using five iterations (ii5). In preparing this manuscript, we also contributed code to the **ALDEx2** package to make the iilr transformation available by providing the argument **`test`** = “**`iterative`**” to the aldex function.

### Differential expression analysis

For each dataset, and for each alignment and quantification protocol, we performed differential expression using the **`edgeR`** package (available for the R programming language) [30] and the **ALDEx2** package [12]. For the simulated data, we evaluated the performance of all differential expression methods using transcript-level abundances. For the real data, we also used gene-level abundances.

When applying **ALDEx2** to the simulated data, we performed DE analysis with each combination of parameters: non-filter versus filter (i.e., the removal of transcripts without at least 10 counts in at least 20 samples), 8 versus 128 Monte Carlo instances, and clr versus iqlr versus malr versus iilr transformation. Immediately, it became obvious that the removal low counts impaired **ALDEx2** precision (*p* < 0.01) for the low variance data. We also observed a non-significant trend in favor of the default recommended 128 MC instances. For clarity of visualization, we prepared figures using only results from “non-filter” runs with 128 Monte Carlo instances. When applying **ALDEx2** to the real data, we used the “non-filter” procedure with 128 Monte Carlo instances. For **ALDEx2**, we considered an expected Benjamini-Hochberg (FDR) adjusted p-value of the Wilcoxon Rank Sum test (i.e., column “wi.eBH") less than 0.05 significant.

When applying **`edgeR`** to the simulated data, we performed DE analysis by applying these functions in order: **`calcNormFactors, estimateCommonDisp, estimateTagwiseDisp`**, and **`exactTest`**. When applying **`edgeR`** to Williams data, we used the **`edgeR`** protocol used in Williams et al. [43]. For **`edgeR`**, we considered an FDR-adjusted p-value of the exact test less than 0.05 significant.

### Performance estimates

For the simulated data, we calculated precision and recall from a contingency table of the simulated state of differential expression (as a binary) compared with the predicted state of differential expression (as a binary). For the real data, consistent with the original publication, we calculated precision and recall for each of the four microarray “truth sets” (available from the supplemental materials of [43]) separately, then reported the average precision and average recall [43].

Since the microarray “truth sets” were based on HGNC symbols, we needed to convert the aligned and quantified transcript-level and gene-level counts to HGNC-level counts; for this, we used code adapted from the Williams et al. methods in conjunction with a conversion table provided by the authors [43]. Using microarray “truth sets” also required an additional filter step to remove HGNC symbols detected by the microarray platform but not RNA-Seq (and *vice versa*). For this, we referenced the **`hgu133plus2.db`**, **`illuminaHumanv4.db`**, and **`illuminaHumanv2.db`** databases (from Bioconductor) to build an HGNC-level “gene universe” for each microarray platform [24]. We then performed a simple set intersection between the microarray “gene universe” with the RNA-Seq HGNC-level “gene universe” prior to calculating precision and recall. Note that our “gene universes” likely differ from those used by Williams et al. (which are not available from their supplemental materials) [43].

## Results

### Benchmark using simulated data

In order to evaluate the performance of **ALDEx2** as a differential expression (DE) method for RNA-Seq data, we tested its performance on three data sets using several combinations of run-time parameters. For the simulated data, we assessed how changes in the alignment and quantification process, sample size, and log-ratio transformation affect the precision and recall of DE analysis. In each comparison, we also performed a DE analysis using edgeR to provide an external point of reference.

### ALDEx2 performance on a simulated data set

Figure 1 shows the precision (top panel) and recall (bottom panel) for a DE analysis of the low variance simulated data set as plotted as a function of software method and log-ratio transformation. Compared with edgeR, clr-based and iqlr-based **ALDEx2** performs just as well in terms of precision. Interestingly, we see that a single iteration of the iilr transformation (i.e., ii1) causes **ALDEx2** to outperform **`edgeR`** in terms of precision. However, regardless of transformation, the recall is very poor with 10 samples (i.e., 5 per group) and still impaired with 20 samples (i.e., 10 per group). Figure 2 shows another projection of these data, plotted against the alignment and quantification procedure. Here, it becomes clear that the choice between **`STAR`** and **`Salmon`** alignment has no apparent impact on the results of DE analysis. Note that Figures 1 and 2 show results from a transcript-level, not gene-level, analysis.

**Figure 1:**
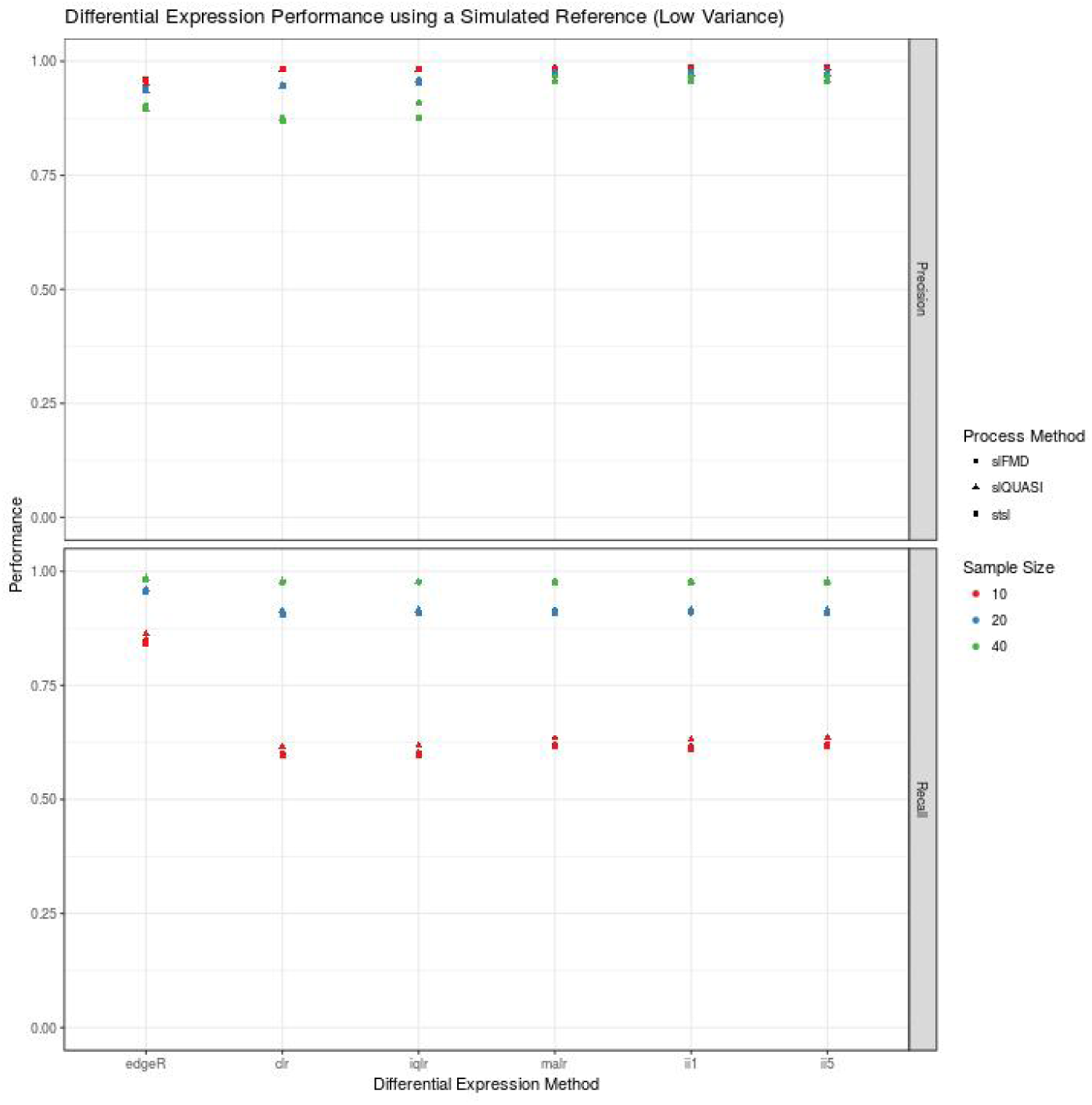
Differential expression analysis of low-variance simulated data. This figure shows the performance (y-axis) of a complete differential expression analysis, organized by differential expression method (x-axis). The acronyms clr, iqlr, malr, ii1, and ii5 describe log-ratio transformations (see Methods). The acronyms slFMD, slQUASI, and stsl describe alignment and quantification procedures (see Methods). Precision (top-panel) is high for **ALDEx2** regardless of log-ratio transformation used. However, **`edgeR`** has much better recall (bottom-panel) than **ALDEx2** for small sample sizes.

**Figure 2:**
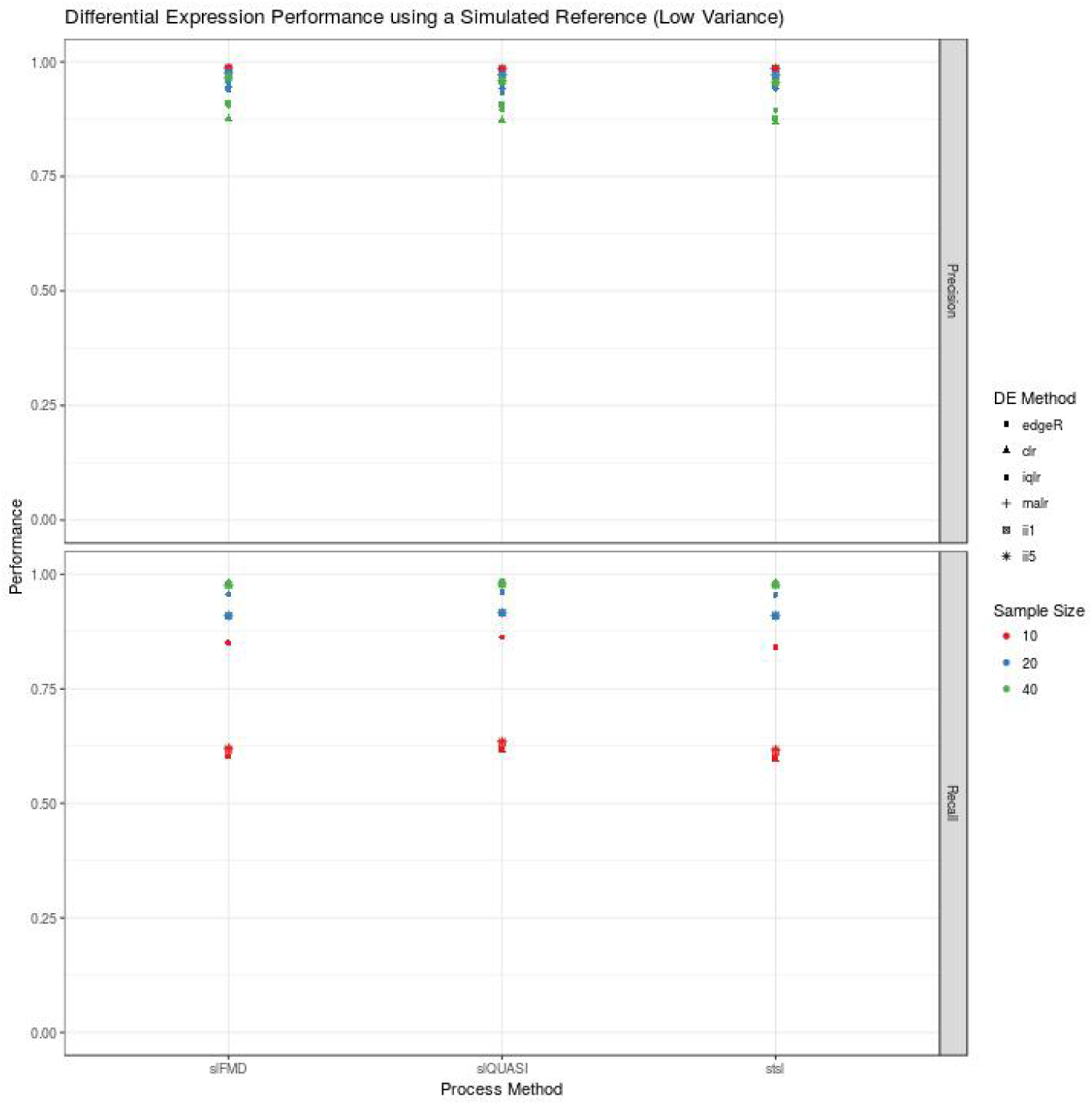
Differential expression analysis of low-variance simulated data. This figure shows the performance (y-axis) of a complete differential expression analysis, organized by alignment and quantification procedure (x-axis). The acronyms clr, iqlr, malr, ii1, and ii5 describe log-ratio transformations (see Methods). The acronyms slFMD, slQUASI, and stsl describe alignment and quantification procedures (see Methods). Precision (top-panel) and recall (bottom-panel) appear largely unaffected by choice in the alignment and quantification procedure.

Figure 3 reproduces Figure 1 for a DE analysis of the high variance simulated data. Here, precision remains high regardless of the approach taken to identify DE transcripts. Recall rates less than 0.25 (even with 40 samples) suggest that the data set is extremely variable. Still, **`edgeR`** clearly outperforms **ALDEx2** in terms of recall. Implicit in this figure is evidence that, again, the choice between **`STAR`** and **`Salmon`** alignment has no apparent impact on the results of DE analysis. Note that Figure 3 shows results from a transcript-level, not gene-level, analysis.

**Figure 3:**
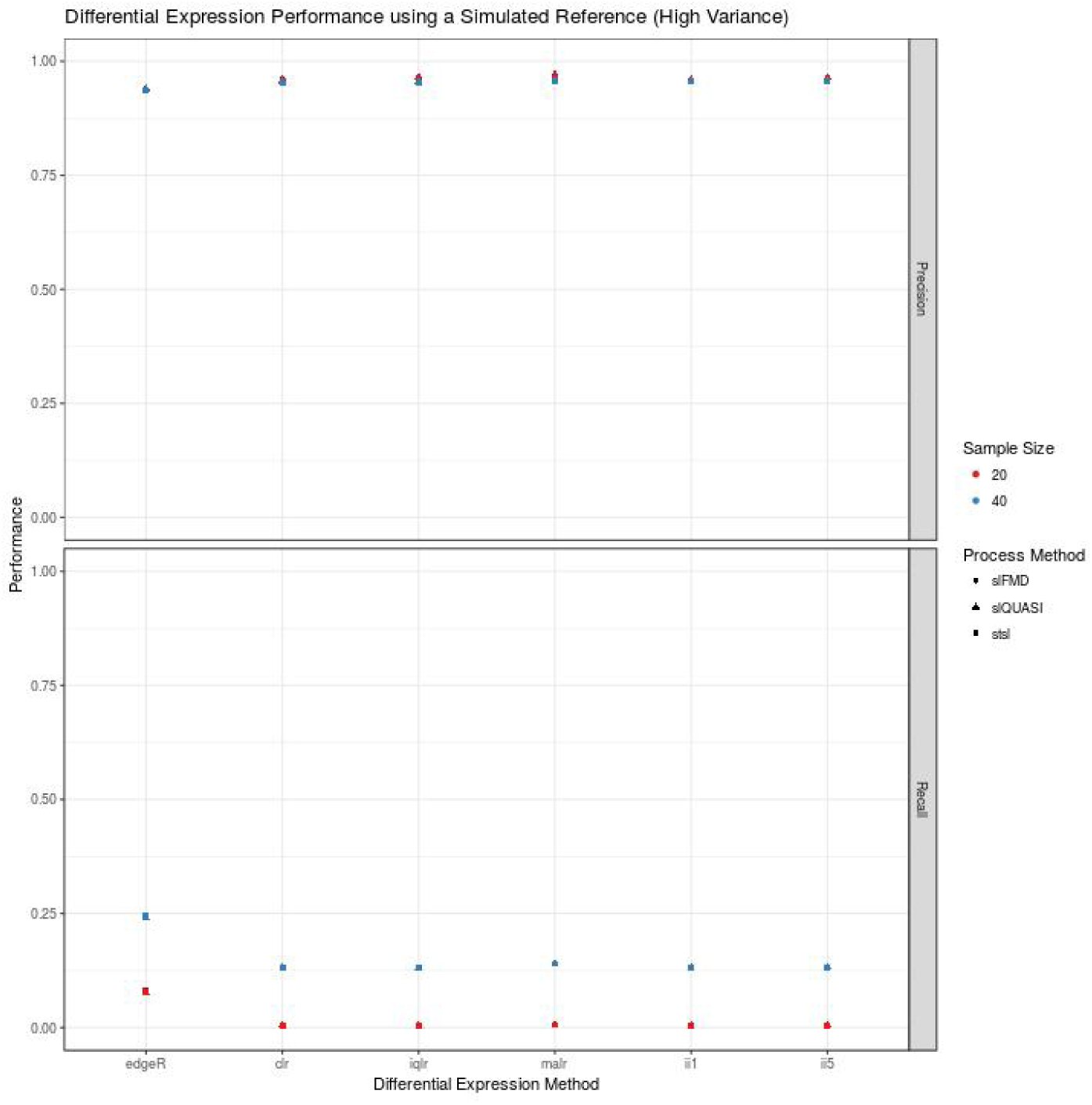
Differential expression analysis of high-variance simulated data. This figure shows the performance (y-axis) of a complete differential expression analysis, organized by differential expression method (x-axis). The acronyms clr, iqlr, malr, ii1, and ii5 describe log-ratio transformations (see Methods). The acronyms slFMD, slQUASI, and stsl describe alignment and quantification procedures (see Methods). Precision (top-panel) is high for **ALDEx2** regardless of log-ratio transformation used. Recall (bottom-panel) is poor for all differential expression method. Again, **`edgeR`** outperforms **ALDEx2** in terms of recall.

### ALDEx2 performance on a real data set

Figure 4 shows precision (y-axis) versus recall (x-axis) for a gene-level (top panel) and transcript-level (bottom panel) DE analysis of the Williams et al. RNA-Seq data (that uses pooled microarray data as a “truth set”) [43]. Here, the smear of lightly colored transparent points indicate the precision and recall recorded by the original Williams et al. publication (sourced from their supplemental materials) [43]. Meanwhile, the dark opaque dots indicate the precision and recall measured during our replication of their procedure. Compared with a myriad of other alignment, quantification, and DE method combinations, **ALDEx2** performs as well as other methods. Interestingly, differences between the choice of log-ratio transformation appear unimportant for these data. Note that the multiplicity of points for each DE method represent different alignment and quantification procedures.

**Figure 4:**
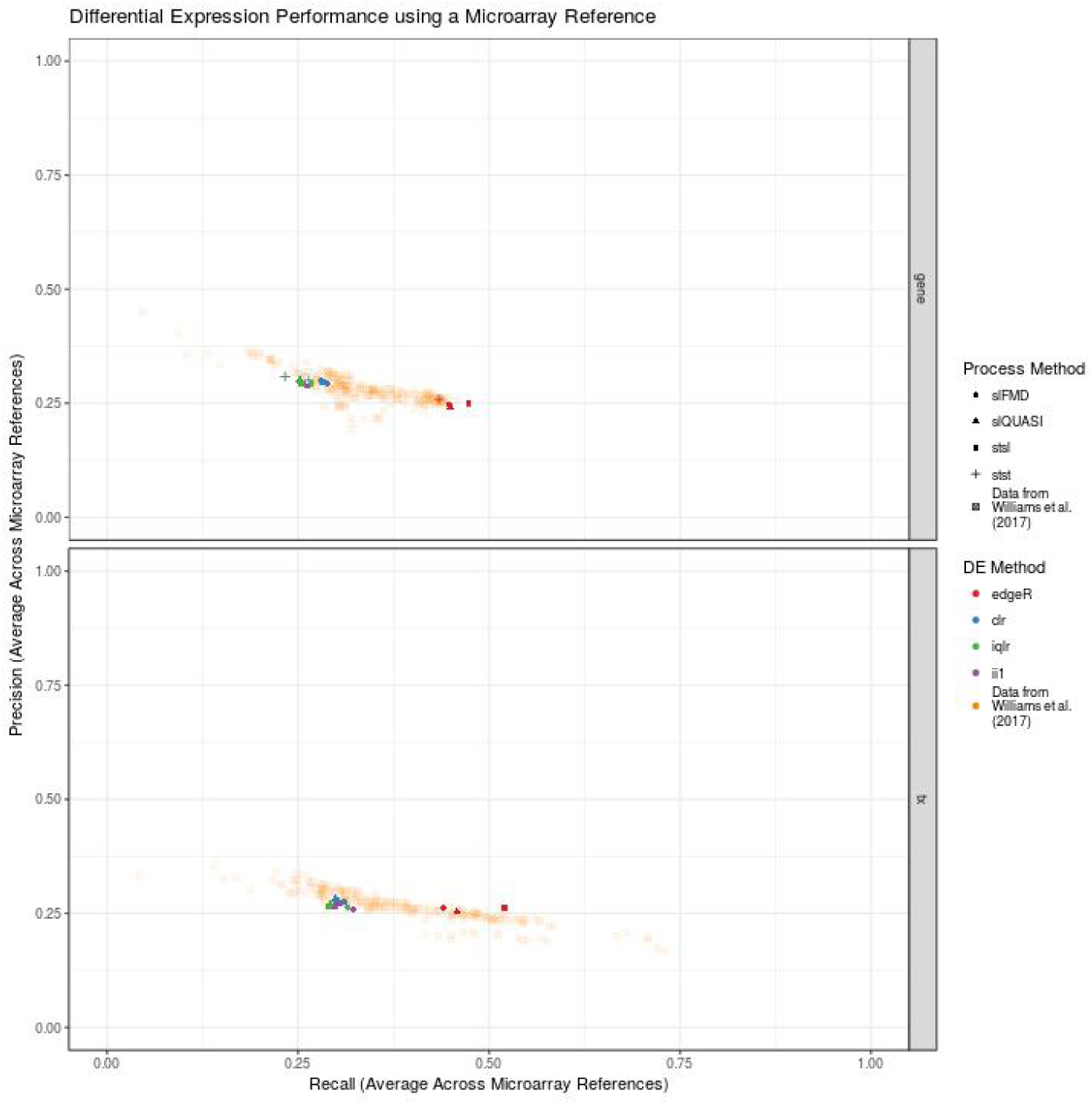
Differential expression analysis of real data. This figure shows the precision (y-axis) versus the recall (x-axis) of a complete differential expression analysis applied to real RNA-Seq data. The “truth set” is established using a microarray reference (see Methods). The acronyms clr, iqlr, and ii1 describe logratio transformations (see Methods). The acronyms slFMD, slQUASI, stsl, and stst describe alignment and quantification procedures (see Methods). Translucent data points show performance calculated from a previously published systematic benchmark. **ALDEx2**, regardless of log-ratio transformation, performs as well as most methods for gene (top-panel) and transcript (bottom-panel) level analyses. Again, **`edgeR`** outperforms **ALDEx2** in terms of recall.

## Discussion

### ALDEx2 has high precision and variable recall for RNA-Seq data

The **ALDEx2** package, most often used to detect differential abundance in 16S rRNA data, has received extensive use for that purpose (e.g., [38] [5]). In previous studies, **ALDEx2** was shown to produce low false discovery rates (FDR) for highly sparse compositional data [36] (FDR =1 – precision). However, we have not yet encountered a study that independently evaluates **ALDEx2** as a differential expression (DE) analysis method for RNA-Seq data (excepting a manuscript by the **ALDEx2** authors which defended its use for RNA-Seq data [13]).

In general, our analysis of simulated and real data agrees with Fernandes et al. [13]: **ALDEx2** can accurately identify differentially expressed genes (and transcripts) in RNA-Seq data. Specifically, **ALDEx2** identifies differentially expressed genes with high precision (i.e., few false positives), but can suffer from low recall (i.e., many false negatives) in the setting of small sample sizes. We offer two explanations for the low recall. First, **ALDEx2** uses non-parametric statistical modeling which tend to have reduced power for small RNA-Seq studies [32] [43] (though, the package authors note that log-ratio transformed data do not necessarily adhere to a normal distribution [13]). Second, methods like edgeR use an empiric Bayes method that “shares information between genes” to shrink per-gene variance estimates and improve power [30]. Presumably, **ALDEx2** would perform better if one could extend moderation to its transformation-based analysis.

Throughout this benchmarking exercise, we had the opportunity to see how two run-time parameters affect **ALDEx2** performance. First, we noted that the removal of lowly abundant counts reduces the precision of **ALDEx2** (*p* < 0.01) (see Supplemental Tables). We suspect that this is because **ALDEx2** can correctly identify differentially expressed transcripts that are all near zero in one of the two groups. Second, we noted that **ALDEx2** performs almost as well using only 8 Monte Carlo instances when compared with using 128 Monte Carlo instances (see Supplemental Tables). Although the package vignette recommends “128 or more mc.samples for the t-test”, this change decreases run-time 16-fold.

### ALDEx2 performance does not depend on alignment and quantification used

The several alignment and quantification procedures used did not change the overall performance of differential expression analysis for **`edgeR`** or **ALDEx2** (regardless of the log-ratio transformation used). This even holds true for the Salmon “quasi-mapping” method that runs many-fold times faster than other quantification algorithms [35]. Although the computational basis of “quasi-mapping” differs from other approaches, this method produces (pseudo-)counts that appear to work well for trimmed M of means (TMM) normalization (used by **`edgeR`**) and log-ratio transformation (used by **ALDEx2**). Broadly speaking, our results agree with the Williams et al. paper in that the choice in differential expression method matters more than the choice in the alignment and quantification method [43].

### ALDEx2 performance depends on log-ratio transformation used

First of all, it is necessary to emphasize that, although log-ratio transformations can be used in lieu of normalization, such transformations do not formally reclaim absolute abundances from relative abundances (see [27]). Yet, benchmarking a transformation-based analysis against a “truth set” implies that the transformation is interpreted as if it were a normalization (i.e., that the reference denominator used for the transformation has rescaled the data to absolute terms [27]). In other words, the more that the reference approximates a feature with fixed abundance across all samples, the more that the transformed data resemble the absolute data. Therefore, the benchmarked performance of a log-ratio transformation-based analysis depends on whether the reference denominator of the transformation is an ideal reference with fixed abundance.

When interpreting the clr transformation as if it were a normalization, there is an implicit assumption that the majority of genes (or transcripts) are not differentially expressed [11]. Meanwhile, the iqlr transformation would assume that a portion of genes (i.e., those with their variances within the inter-quartile range) are not differentially expressed. Likewise, the iterative transformation assumes that the results of an **ALDEx2** analysis selects (non-)differentially expressed genes more accurately than a simpler transformation. Therefore, all else being equal, one can interpret the performance of each transformation as a proxy for how well it reclaims absolute information (i.e., how well it approximates an ideal reference with fixed abundance).

For simulated data, some transformations perform better than others. First, the iqlr transformation is more precise than the clr transformation. This is expected because the iqlr transformation should be more robust to imbalances in up- or down-regulation [44]. Second, the iterative transformations is more precise than the iqlr, suggesting that novel log-ratio transformations could improve performance beyond those routinely used. Perhaps surprisingly, this trend was not apparent among the real data. This could be due to either the absence of any imbalance in this data set (such that iqlr and others offer no clear benefit over clr) or limitations in the microarray-based “truth set” (which may not have controlled for a systematic imbalance in the biological samples).

### Limitations of ALDEx2 and other transformation-based methods

Users may encounter some practical limitations with **ALDEx2** as it is currently implemented. First, owing to the replication of non-parametric analyses across multiple Monte Carlo instances, **ALDEx2** runs much slower than edgeR and other methods (though in preparing this manuscript we submitted code to speed up the software). Second, **ALDEx2** does not contain a generalization to mixed models. Although this package does have an ANOVA-like procedure (i.e., using Kruskal-Wallis), it takes even longer to run than the standard two-group comparisons.

There are also limitations that come with interpreting the log-ratio transformation as a kind of normalization. As mentioned above, using the clr in this way assumes that the majority of genes (or transcripts) are not differentially expressed [11], the same assumption held by the trimmed mean of M (TMM) normalization [31]. When this assumption does not hold, it is not possible to infer absolute differences in expression between samples. As such, both will fail if one of the experimental groups has massively more up-regulation than down-regulation (or *vice versa*). This scenario is exemplified by a cell line with high levels of c-Myc, a protein with widely variable expression in tumors that can amplify gene expression to produce 2-3 times more total RNA [21]. When comparing high c-Myc cells against low c-Myc cells, a standard RNA-Seq analysis would conclude that there is both up- and down-regulation, even though the cells actually only have up-regulation (i.e., based on using spike-in controls normalized to cell number) [21]. A transformation-based approach would not fair any better here, unless one was exclusively interested in identifying transcripts that were differentially expressed *relative to a reference* [27].

Finally, **ALDEx2** may suffer from another limitation based on how it uses the Dirichlet distribution to generate Monte Carlo instances. Weiss et al. note, “this formulation assumes a Dirichlet-multinomial framework, which imposes a negative correlation structure on every pair of [features]” [42]. In fact, Hawinkel et al. showed that, when simulating 16S abundance data to have a positive correlation structure, **ALDEx2** results depart from the uniformity of the *p*-value distribution (in the liberal direction) and show an increase in the nominal false discovery rate (FDR) (although FDR-adjustment still brought FDR below the 0.05 threshold) [17]. However, this publication also found that a positive correlation structure negatively impacted the performance of popular differential expression analysis methods like **DESeq2** and **edgeR** (the latter of which had very high FDR-adjusted FDR rates) [17].

## Conclusions

Across all applications, the **ALDEx2** package has high precision in identifying differentially expressed genes (and transcripts) from simulated and real RNA-Seq data. With sufficient sample sizes, it also has good recall. Across all log-ratio transformations, the removal of lowly abundant counts before differential expression analysis impedes **ALDEx2** performance. Meanwhile, the choice in alignment and quantification procedure does not seem to affect the results of an **ALDEx2** analysis. However, at least when interpreting log-ratio transformations as if they were normalizations, some transformations may work better than others. Nevertheless, our results suggest that log-ratio transformation-based methods can work to measure differential expression from sequencing data, provided that certain assumptions are met.

## Additional Files

### Additional file 1 — Analysis scripts

This file contains all the scripts used to generate the simulated data, benchmark methods on the simulated data, benchmark methods on the real data, parse the results, and make the figures.

### Additional file 2 — Simulated data (low variance) benchmark performance

This table contains the precision and recall estimates for several methods as applied to the low variance simulated data. This table is used to make figures.

### Additional file 3 — Simulated data (high variance) benchmark performance

This table contains the precision and recall estimates for several methods as applied to the high variance simulated data. This table is used to make figures.

### Additional file 4 — Real data benchmark performance

This table contains the precision and recall estimates for several methods as applied to the real data. This table is used to make figures.

